# Cardiomyocyte Contractile Impairment in Heart Failure Results from Reduced BAG3-mediated Sarcomeric Protein Turnover

**DOI:** 10.1101/2020.04.10.022319

**Authors:** Thomas G. Martin, Valerie D. Myers, Praveen Dubey, Shubham Dubey, Edith Perez, Christine S. Moravec, Monte S. Willis, Arthur M. Feldman, Jonathan A. Kirk

**Affiliations:** Department of Cell and Molecular Physiology, Loyola University Stritch School of Medicine, Maywood, IL; Department of Medicine, Temple University Lewis Katz School of Medicine, Philadelphia, PA; Department of Medicine, Cleveland Clinic Lerner College of Medicine, Cleveland, OH; Department of Pathology and Laboratory Medicine, Indiana University School of Medicine, Indianapolis, IN

## Abstract

The association between reduced myofilament force-generating capacity (F_max_) and heart failure (HF) is clear, however the underlying molecular mechanisms are poorly understood. Here, we show the F_max_ decrease arises from impaired BAG3-mediated sarcomere turnover. Myofilament BAG3 decreased in human HF and predicted the extent of F_max_ decrease. This relationship was confirmed using BAG3^+/-^ mice, which had reduced F_max_ and increased myofilament ubiquitination, suggesting impaired protein turnover. We show cardiac BAG3 operates via the chaperone-assisted selective autophagy complex (CASA), conserved from skeletal muscle, and confirm sarcomeric CASA localization is BAG3/proteotoxic stress-dependent. To determine if increasing BAG3 expression in HF would restore sarcomere proteostasis/F_max_, HF mice were treated with AAV9/BAG3. Gene therapy fully restored F_max_ after four weeks and decreased ubiquitination. Using mass spectrometry, we identified several sarcomere proteins with increased ubiquitination in HF and four that decreased with AAV9/BAG3. Our findings indicate BAG3-mediated sarcomere turnover is required for myofilament functional maintenance.

## INTRODUCTION

Heart failure is the leading cause of morbidity and mortality in the industrialized world and is characterized by impaired contractility and decreased cardiac output^1^. At the cellular level, several factors have been implicated in the development of heart failure, including defective calcium handling, neurohormonal imbalance, and functional decline of the sarcomere – the fundamental molecular unit of contraction^2–4^. The sarcomere is a paracrystalline protein complex that mediates cellular contraction through myosin and actin filaments, which engage in the calcium-dependent cross-bridge cycle. Increased myofilament calcium sensitivity and decreased maximum force-generating capacity (F_max_) are well-documented in heart failure^5–10^. The altered calcium sensitivity is uniformly attributed to changes in site-specific post-translational modifications of myofilament proteins, most commonly phosphorylation of troponin I^11^. However, the molecular mechanisms for decreased F_max_ in heart failure are incompletely understood.

The sarcomere is under constant mechanical stress, which predisposes its members to denaturation, and yet must maintain optimal mechanical function for decades in the terminally differentiated adult cardiomyocyte. To preserve function, stress-denatured sarcomere proteins must be refolded or targeted to degradation pathways by molecular chaperones so that new proteins can be incorporated^12^. However, sarcomeric protein quality control is frequently impaired in heart failure, leading to aggregation of misfolded/dysfunctional proteins^12^. An elevation of dysfunctional proteins incorporated into the sarcomere in heart failure may thus serve as a plausible mechanism for the observed decrease in F_max_. However, the distinct mechanisms of sarcomere protein turnover in the adult heart are poorly characterized, and the myofilament functional relevance of sarcomere protein quality control has not been determined.

Bcl2-associated athanogene 3 (BAG3) is a heat shock protein co-chaperone involved in protein degradation through both chaperone-mediated autophagy and selective macroautophagy^13^. Evidence in recent years suggests a role for BAG3 in sarcomere structural and functional maintenance. Numerous clinical studies identified BAG3 mutations and decreased BAG3 expression are associated with dilated cardiomyopathy (DCM) and myofibrillar myopathy in humans^14–20^. Due to difficulties with maintaining adult cardiomyocytes in culture, mechanistic studies of BAG3 were primarily performed in immature neonatal myocytes, where BAG3 was shown to stabilize sarcomere structure through maintenance of the actin capping protein CapZβ^21^. However, the role of BAG3 at the sarcomere in adult myocytes under mechanical load is less understood and the association with CapZβ does not appear to be conserved^22^. Moreover, germline KO of BAG3 does not impact sarcomere structure in neonatal myocytes, which rapidly disintegrates post-natum^23^. These findings suggest separate or additional sarcomeric roles for BAG3 in the adult cardiomyocyte.

An important *in vivo* study of BAG3 in the adult heart came from Fang *et al*. who showed BAG3 KO in mice caused DCM, destabilized the small HSPs (HSPBs), and resulted insoluble protein aggregation, indicating impaired protein turnover^24^. Interestingly, BAG3 KO in iPSC-derived cardiomyocytes resulted in decreased myocyte contractility and altered expression of HSPs, suggesting potential sarcomeric functional relevance for BAG3-dependent protein quality control^25^. However, iPSC-cardiomyocytes also have immature morphology and are not under mechanical load, making functional inferences to the adult problematic, especially at the sarcomere, which often matures only under chronic load. While little is known regarding BAG3’s role in the mature cardiac sarcomere, studies in mature skeletal muscle identified BAG3 assembles with HSP70 and HSPB8, forming the chaperone-assisted selective autophagy (CASA) complex which mediates the degradation of mechanically misfolded filamin C^26–28^. Whether CASA operates at the cardiac sarcomere, the myofilament functional impact of BAG3, and the effect of heart failure on these mechanisms are unknown. However, given the near universally observed depression of F_max_ in systolic heart failure, sarcomeric protein quality control represents an attractive therapeutic target.

Here, we identify sarcomere protein turnover is impaired in human DCM samples with reduced F_max_. We confirm the presence of the CASA complex at the cardiac sarcomere and show myofilament BAG3 expression decreases in DCM and is directly associated with functional decline, where lowest BAG3 expression predicted weakest force generation. The functional significance of BAG3 at the sarcomere is further exemplified in BAG3^+/-^ mice, which display reduced F_max_ and increased myofilament ubiquitination. These mice also have reduced expression of several CASA proteins in the myofilament fraction, suggesting BAG3-dependent localization. Using a mouse heart failure model with BAG3 gene therapy, we identify that increasing BAG3 expression in heart failure restores myofilament F_max_ and sarcomere proteostasis. Finally, using ubiquitin-enrichment mass spectrometry we show BAG3 gene therapy restores turnover of four sarcomere proteins: the previously identified CASA client filamin C, and three new clients – tropomyosin, sarcomeric α-actin, and clathrin heavy chain. Our novel findings indicate impaired sarcomere protein turnover contributes to decreased F_max_ in heart failure and highlight BAG3 as an essential factor for sarcomere functional maintenance.

## RESULTS

### Human DCM cardiomyocytes have reduced sarcomere contractile function and increased myofilament protein ubiquitination

Decreased contractility in heart failure often stems from sarcomere dysfunction^3^. However, the mechanisms underlying reduced sarcomere force generation (F_max_) in heart failure remain to be fully characterized. In genetic cases of heart failure, decreased F_max_ may be explained by structural alterations to proteins involved in the cross-bridge cycle, which are detrimental to their function^29,30^. However, functional deficits have also been observed in non-genetic heart failure^6,7,9,29^, which makes up the majority of cases. We therefore sought to determine the underlying mechanism for myofilament dysfunction in heart failure using left ventricle (LV) samples from patients with idiopathic dilated cardiomyopathy (DCM) (**Table 1**).

**Table 1.** Clinical characteristics of the human DCM and non-failing samples.

| **Characteristic** | **NF (n = 9)** | **DCM (n = 21)** | **P value** |
| --- | --- | --- | --- |
| Age, (years), Mean ± SD | 53.3 ± 8.9 | 55.5 ± 11.5 | 0.66 |
| Female, n, (%) | 4 (44.4) | 8 (38.1) |  |
| Ethnicity, n, (%) |  |  |  |
| - White | 8 (88.9) | 16 (76.2) |  |
| - Black | 1 (11.1) | 4 (19.1) |  |
| - Hispanic | 0 (0.0) | 1 (4.8) |  |
| EF, (%), Mean ± SD | 57.5 ± 3.2 | 20.3 ± 2.2 | < 0.001 |

Using skinned myocytes from human DCM and non-failing (NF) samples, we first showed that myofilament F_max_ decreased by ~45% in DCM (**Figure 1A-B**). Consistent with prior studies discussed above, the EC_50_ (calcium concentration required to elicit half maximal force) also decreased in DCM compared with NF, indicating increased myofilament calcium sensitivity (**Figure 1C**). Since impairment to protein quality control mechanisms is a common feature of heart failure^31^, we hypothesized that decreased turnover of sarcomere proteins might underly the observed decrease in force-generation. To determine if sarcomere proteostasis was impaired, we assessed protein ubiquitination in myofilament-enriched fractions from the DCM and NF tissue by western blot. As expected, we found significantly increased ubiquitin in DCM, suggesting inadequate turnover of sarcomere proteins in disease (**Figure 1D-E**).

**Figure 1.**
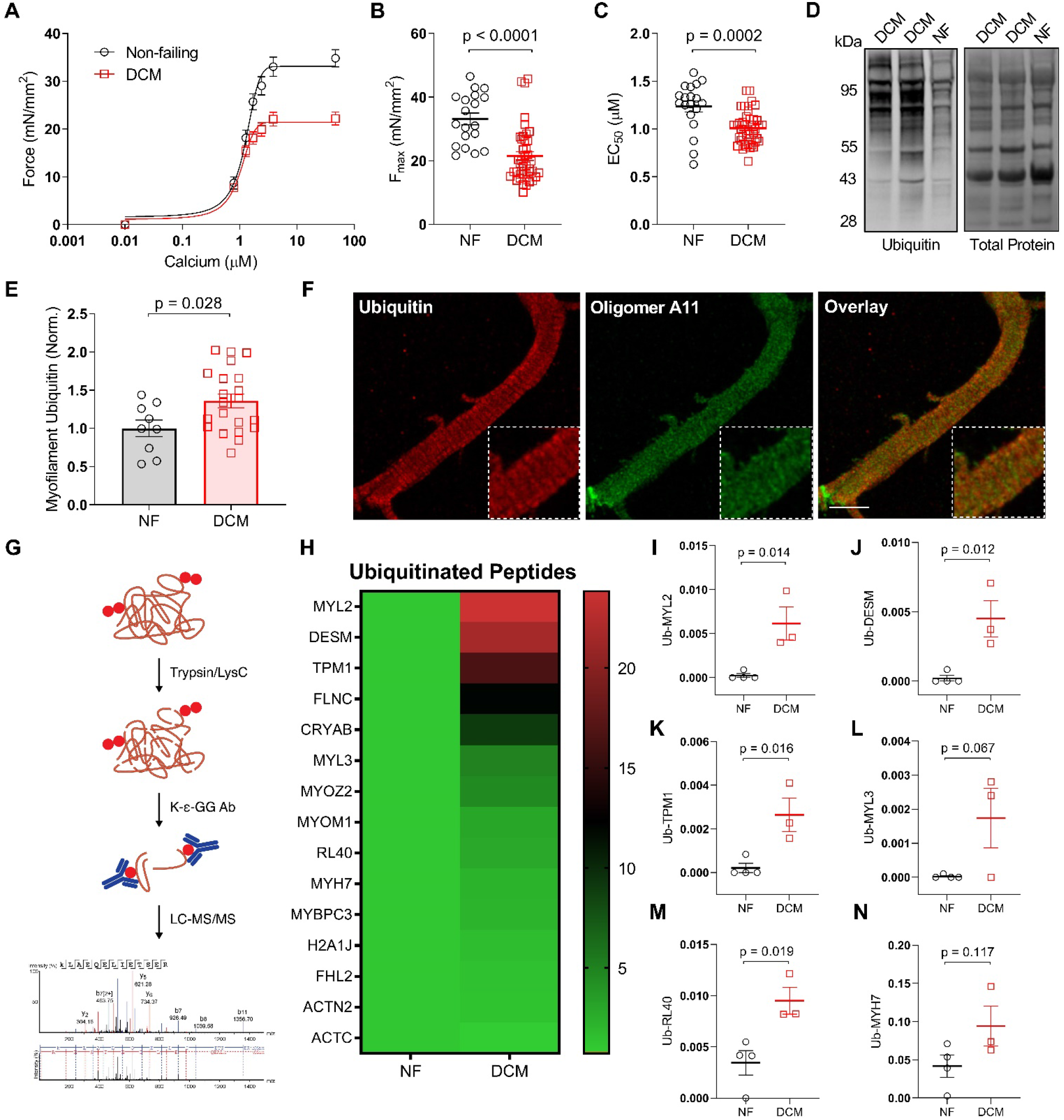
Human DCM is characterized by decreased myofilament function and impaired sarcomere protein turnover. **A.** Skinned myocyte force-calcium relationship from human NF and DCM cardiomyocytes. **B-C.** Summary data for myocyte F_max_ (B) and EC_50_ (C); n = 19 NF from 6 patients, 41 DCM from 12 patients. **D-E.** Representative western blot for myofilament ubiquitin (D) and normalization to total protein loading control (E) in NF and DCM samples; n = 9 NF, 21 DCM. **F.** Representative immunofluorescence image of a human LV cardiomyocyte immunostained for ubiquitin and oligomer A11; 63X magnification, scale bar = 10 μm. **G.** Simplified proteomics paradigm for ubiquitinated peptide enrichment. **H.** Heat map of ubiquitinated proteins identified by LC-MS/MS for NF and DCM human patients; scale bar = fold change relative to NF. **I-N.** Spectral count data for ubiquitinated peptides normalized to total peptide input for myosin regulatory light chain 2 (I), desmin (J), tropomyosin (K), myosin light chain 3 (L), ribosomal protein L40 (M), and β-MHC (N); n = 4 NF, 3 DCM. All data are presented as mean ± SEM and were analyzed by 2-tailed Student’s *t*-test.

There are two possible explanations for this observed increase in myofilament ubiquitination. One is that these ubiquitinated proteins were removed from the contractile apparatus, but not degraded properly and thus formed protein aggregates that are commonly observed in heart failure^31^. These aggregates are associated with contractile dysfunction^31^, providing a possible explanation for the decrease in F_max_. However, a second possibility, which has been suggested in the skeletal muscle sarcomere during atrophy^32^, is that old/misfolded sarcomeric proteins are not being removed from the contractile apparatus and remain integrated into the sarcomere. The protein stoichiometry of the sarcomere is highly conserved^33^, and thus the continued integration of these old sarcomeric proteins which have been marked for degradation would block the introduction of newly synthesized proteins, providing an explanation for a decline in F_max_. This second mechanism is supported by the increased protein ubiquitination detected in the myofilament-enriched fraction (**Figure 1D, E**).

To determine whether these ubiquitinated proteins are embedded within the sarcomere itself rather than in protein aggregates, we used super-resolution confocal microscopy on skinned human DCM cardiomyocytes immunostained for ubiquitin and the protein aggregate marker oligomer A11. As expected, A11 had diffuse punctate localization with no apparent sarcomere patterning. However, ubiquitin localized in a cross-striated pattern and was enriched in thick, regularly repeating bands representative of the sarcomere A-band (**Figure 1F**). Among the most abundant A-band proteins are β-MHC, actin, and myosin binding protein C, which are fundamental to sarcomere contraction^34^. This indicates the ubiquitinated proteins identified in the myofilament fraction, which were found to increase in DCM by western blot, are in large part incorporated into the sarcomere and not components of protein aggregates. It further suggests the primary proteins with inadequate turnover in DCM are those most essential to myofilament force generation.

To determine which specific sarcomere proteins had impaired turnover, we digested proteins from the myofilament fraction isolated from NF and DCM tissue with Trypsin/LysC protease, enriched for ubiquitinated peptides by immunoprecipitation for the diglycine lysine ubiquitin remnant motif, and identified the peptides using liquid chromatography tandem mass spectrometry (LC-MS/MS) (**Figure 1G**). To our knowledge, this is the first use of ubiquitin-enrichment mass spectrometry in the human heart. Moreover, this technique provided the first ever characterization of the myofilament ubiquitinome in heart failure and identified several proteins with impaired turnover. The top hits with increased ubiquitination in DCM included myosin regulatory light chain 2 (MLRV), desmin (DESM), and tropomyosin alpha 1 (TPM1), which had significantly higher ubiquitination when analyzed by spectral count compared to non-failing (**Figure 1H-N**). Collectively, these data indicate sarcomere protein turnover is disrupted in DCM resulting in increased misfolded proteins incorporated into the sarcomere, which offers a compelling possible explanation for the decrease in F_max_.

### Myofilament expression of the co-chaperone BAG3 decreases in DCM, which is directly associated with decreased F_max_

Several studies have implicated a role for the heat shock protein co-chaperone BAG3 in sarcomere maintenance^20,21,23,35,36^, however, the mechanisms of BAG3-mediated sarcomere maintenance in adult cardiomyocytes are poorly understood. BAG3 is involved in protein quality control by mediating autophagic clearance of misfolded proteins and has been shown to localize to the sarcomere Z-disc, thus making it a potential candidate for mediating sarcomere protein turnover^37,38^. One previous study found whole LV BAG3 expression decreased in end-stage heart failure, however, myofilament BAG3 has not previously been assessed^15^. We used western blot to determine myofilament-localized BAG3 expression in human DCM and NF samples and found BAG3 levels decreased significantly in DCM (**Figure 2A-B**). Strikingly, when we compared myofilament BAG3 levels with sarcomere functional parameters in the DCM samples, F_max_ was positively associated with BAG3 expression where samples with highest BAG3 expression displayed the strongest F_max_ (**Figure 2C**). As expected, there was no association between BAG3 expression and calcium sensitivity in DCM (**Figure 2D**). Importantly, this relationship is specific to the myofilament pool of BAG3, as there was no association between cytosolic/soluble BAG3 and F_max_ (**Figure S1**).

**Figure 2.**
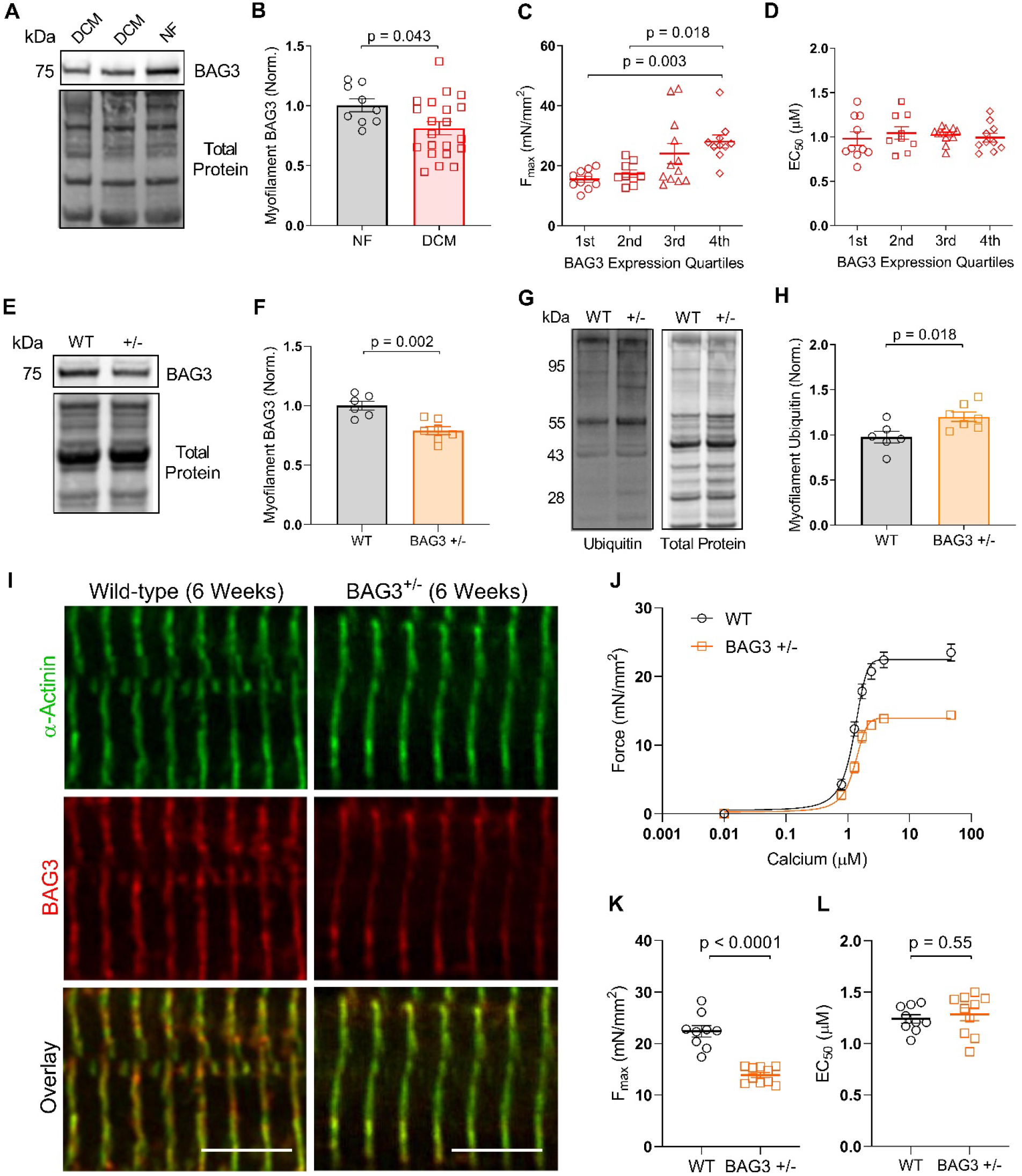
Sarcomeric BAG3 expression decreases in DCM and is associated with reduced myofilament F_max_. **A-B.** Representative western blot for myofilament BAG3 (A) and normalization to total protein loading control (B) in NF and DCM humans; n = 9 NF, 21 DCM; 2-tailed *t*-test. **C-D.** DCM myocyte F_max_ (C) and EC_50_ (D) grouped by quartile of BAG3 expression; 1^st^ = lowest BAG3 expressors; n = 12 DCM samples, 3-4 myocytes/sample for functional assessment; one-way ANOVA, Tukey post-hoc. **E-F.** Representative western blot for myofilament BAG3 (E) and normalization to total protein loading control (F) in WT and BAG3^+/-^ mice. **G-H.** Representative western blot for myofilament ubiquitin (G) and normalization to total protein loading control (H) in WT and BAG3^+/-^ mice; n = 6 WT, 7 BAG3^+/-^; 2-tailed *t*-test. **I.** Representative immunofluorescence image of WT and BAG3^+/-^ cardiomyocytes immunostained for α-actinin and BAG3; 63X magnification, scale bars = 5 μm. **J.** Skinned myocyte force-calcium relationship from WT and BAG3^+/-^ cardiomyocytes. **K-L.** Summary data for myocyte F_max_ (K) and EC_50_ (L); n = 9 WT from 3 mice, 10 BAG3^+/-^ from 3 mice; 2-tailed *t*-test. All data are presented as mean ± SEM.

To further examine the impact of reduced BAG3 expression on myofilament function, we next employed a mouse model with cardiomyocyte-restricted heterozygous BAG3 expression (BAG3^+/-^). These mice were previously found to have progressive LV dysfunction starting at 8 weeks of age^39^. In the present study, we used 6-week-old mice to capture the early effects of BAG3-specific haploinsufficiency on myofilament function, rather than the general response to heart failure development. At 6 weeks, the mice displayed a ~20% reduction in myofilament BAG3 compared to wild-type and had significantly increased myofilament protein ubiquitination (**Figure 2E-H**). Extensive sarcomere structural abnormalities, which have been identified in BAG3 germline KO mice at 3 weeks^23^ and in BAG3 KO iPSC-cardiomyocytes^35^, were not observed in the BAG3^+/-^ mice at this time-point (**Figure 2I**). We next assessed myofilament function using force-calcium experiments on skinned myocytes from the BAG3^+/-^ and WT mice and found cardiomyocytes from BAG3^+/-^ mice had severely reduced F_max_ but no change in calcium sensitivity (**Figure 2J-L**). These data indicate BAG3 is required for optimal contractile function of the sarcomere and for maintaining sarcomere proteostasis. Additionally, sarcomere dysfunction due to BAG3 haploinsufficiency arises despite normal sarcomere morphology, suggesting impaired sarcomere protein turnover – not structural disarray – may be the initial insult that leads to contractile dysfunction and ultimately heart failure. It further agrees with our finding that the ubiquitinated proteins remain integrated in the sarcomere and induce dysfunction but not disarray.

As previously discussed, numerous BAG3 mutations have been linked to heart failure in humans. One particularly penetrant mutation, the missense proline to leucine at amino acid 209 (P209L), precludes ubiquitinated substrate targeting to the autophagy pathway and is associated with protein aggregation, restrictive cardiomyopathy, and severe myofibrillar myopathy^20,36,40^. An earlier study found that mice with cardiomyocyte-restricted expression of the human P209L BAG3 transgene develop restrictive cardiomyopathy by 8 months of age, which is accompanied by elevated ubiquitin and protein aggregation^41^. We found that protein ubiquitination in the myofilament fraction also increases in the P209L mice (**Figure S2**). Therefore, we employed this model to directly assess the impact of disrupted BAG3-mediated sarcomere protein turnover on myofilament function. Using skinned myocyte force-calcium experiments on cardiomyocytes from 8-month-old P209L mice, we found F_max_ was significantly reduced compared with aged-matched littermate controls (**Figure S2**). Taken together, the relationship between decreased/dysfunctional BAG3 and reduced force-generating capacity identified in the human DCM samples, the BAG3^+/-^ mice, and the P209L mice indicates BAG3 is essential for maintaining maximum sarcomere tension generation in the absence of overt structural disarray.

### The chaperone-assisted selective autophagy (CASA) complex localizes to the sarcomere Z-disc and is upregulated in response to proteotoxic stress

Having established that BAG3 is functionally significant for the sarcomere, we next sought to identify the underlying mechanism. To begin, we used immunoprecipitation for myofilament BAG3 in the human LV and identified the BAG3 interactome via mass spectrometry. Mass spectrometry analysis identified numerous sarcomere members, which may represent clients of BAG3-mediated protein turnover, as well as two heat shock proteins: Hsp70 and HspB8 (**Table S1**). These two binding partners are noteworthy, as together with BAG3 they were previously shown to engage in chaperone-assisted selective autophagy (CASA) in skeletal muscle^26,27^. In CASA, Hsp70 and HspB8 associate with BAG3 and bind to mechanically misfolded proteins, which are ubiquitinated by the E3 ligase c-terminus of Hsp70 interacting protein (CHIP) and targeted to the autophagosome for degradation^28^. To confirm the presence of the CASA complex at the cardiac sarcomere, we used immunofluorescence microscopy on human LV cardiomyocytes and found BAG3, Hsp70, HspB8, and CHIP each localized to the sarcomere Z-disc, as evident from their co-localization with α-actinin (**Figure 3A-B**). We further confirmed this localization was sarcomeric using western blot for the CASA members in human whole LV, soluble, and myofilament fractions (**Figure 3C**) and confirmed their association therein by co-immunoprecipitation (**Figure 3D-E**). These data confirm that the CASA complex identified in skeletal muscle is conserved at the sarcomere in human cardiomyocytes.

**Figure 3.**
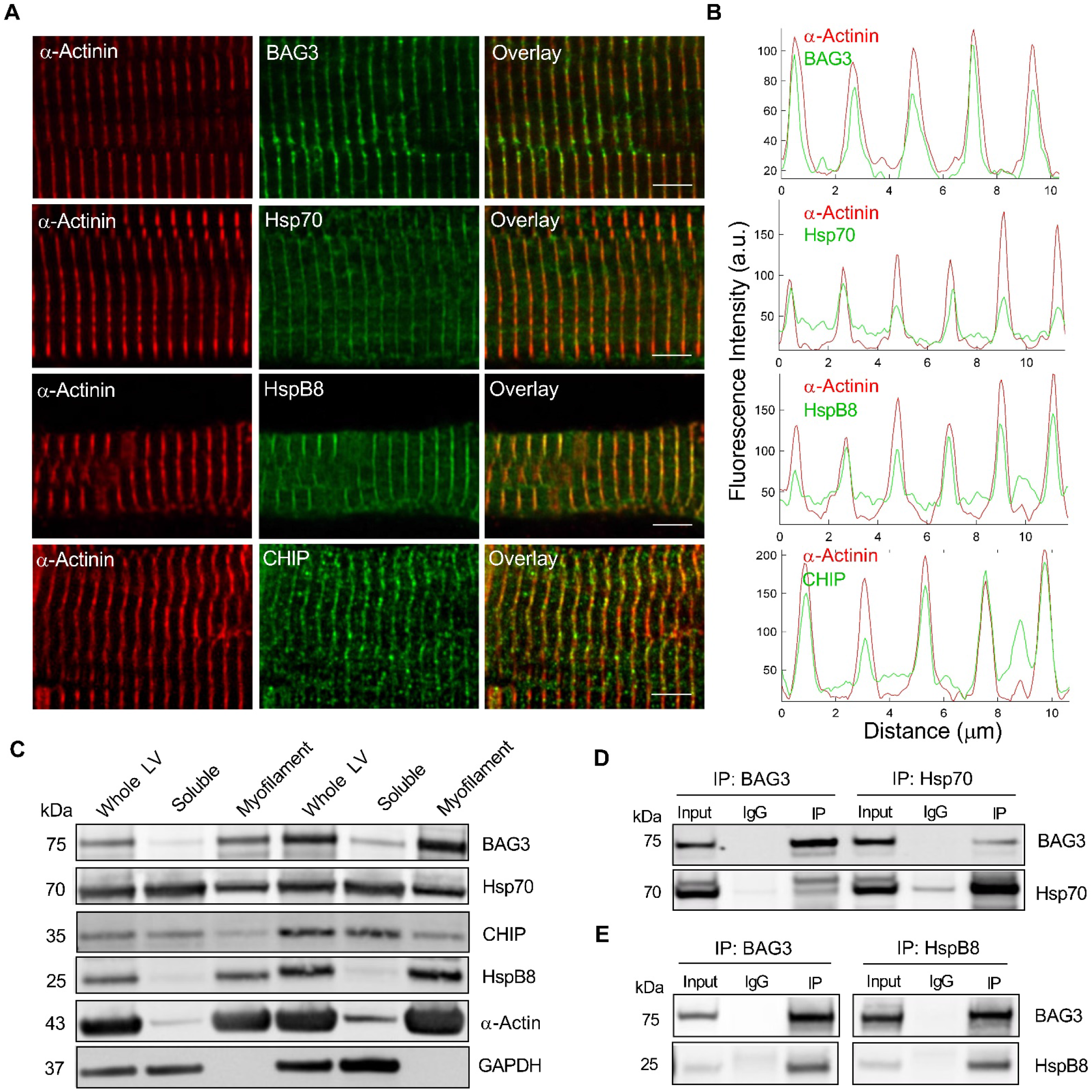
The CASA complex localizes to the sarcomere Z-disc in cardiomyocytes. **A.** Representative immunofluorescence images of human LV cardiomyocytes immunostained for BAG3, Hsp70, HspB8, and CHIP and counterstained for the Z-disc protein α-actinin; 63X magnification, scale bars = 5 μm. **B.** Quantitative line scan of fluorescence intensity from the corresponding green and red channels in A over distance; a.u. = arbitrary units. **C.** Representative western blot for the CASA complex proteins in the whole LV, triton-soluble, and myofilament fractions; sarcomeric α-actin = myofilament fraction positive control, GAPDH = soluble fraction positive control. **D.** Western blot showing results of reciprocal co-immunoprecipitation for BAG3 and Hsp70 in myofilament fraction. **E.** Western blot showing results of reciprocal co-immunoprecipitation for BAG3 and HspB8 in myofilament fraction.

Protein chaperones and co-chaperones are regulated by cell stress, advanced levels of which stimulate increased chaperone protein expression. This was previously shown for BAG3 in a fibroblast-like cell line, where treatment with the proteasome inhibitor MG132 caused an upregulation of BAG3 expression^42^. To determine if the CASA complex exhibits proteotoxic stress-dependent localization to the sarcomere, we isolated neonatal rat ventricular myocytes (NRVMs) from one-day-old rats and maintained them in culture. To cause proteotoxic stress, the NRVMs were treated with 2 μM MG132 for 24 hours. MG132 treatment caused a pronounced upregulation of BAG3, Hsp70, HspB8, and CHIP expression in the myofilament fraction, indicating that CASA does indeed localize to the sarcomere in response to proteotoxic stress (**Figure 4A-G**). This finding was substantiated further by super-resolution confocal imaging of DMSO and MG132-treated NRVMs, which showed a prominent increase of Z-disc localized BAG3 in response to MG132 (**Figure 4H-K**).

**Figure 4.**
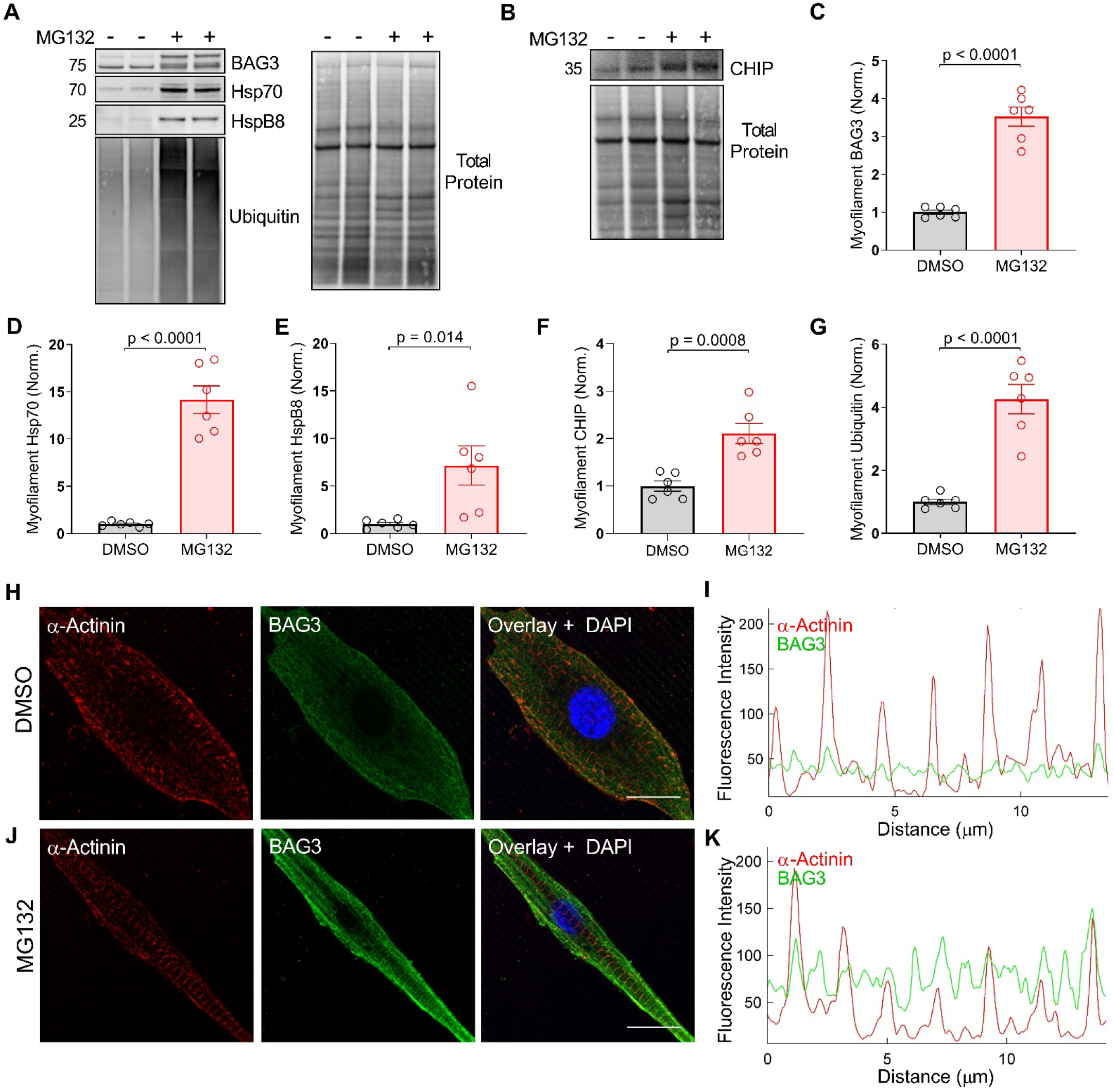
The CASA complex is targeted to the sarcomere in response to proteotoxic stress. **A-B.** Representative western blots for myofilament BAG3, Hsp70, HspB8, ubiquitin, and CHIP in NRVMs treated with DMSO or the proteasome inhibitor MG132. **C-G.** Quantification of myofilament expression normalized to total protein loading control for BAG3 (C), Hsp70 (D), HspB8 (E), CHIP (F), and ubiquitin (G); n = 6 DMSO, 6 MG132 from three separate biological samples; 2-tailed *t*-test. **H-K.** Representative immunofluorescence images for NRVMs treated with DMSO (H) and MG132 (J) immunostained for BAG3 and α-actinin, with quantitative line scan of fluorescence intensity (I, K); 63X magnification, scale bars = 10 μm.

### HspB8 and CHIP display BAG3-dependent association with the myofilament

We next asked whether BAG3 was required for localization of the other CASA members to the myofilament. To test whether the assembly of the CASA complex proteins at the myofilament was BAG3-dependent, we used mouse models of cardiomyocyte-specific heterozygous and homozygous BAG3 deletion. Using western blot in myofilament-enriched LV tissue from wild-type, BAG3^+/-^, and BAG3^-/-^ mice, we found myofilament levels of HspB8 and CHIP were significantly reduced in the partial and complete absence of BAG3 (**Figure 5A-D**). However, myofilament Hsp70 expression was not impacted by decreased BAG3, suggesting it requires different mechanism for targeting to the sarcomere. This finding is not surprising as Hsp70 is a general chaperone for numerous protein clients and operates in many arms of cellular protein folding and degradation^43^. Nevertheless, that myofilament CHIP was reduced with decreased BAG3 is likely significant for all Hsp70-mediated degradation processes as CHIP regulates Hsp70-client ubiquitination.

**Figure 5.**
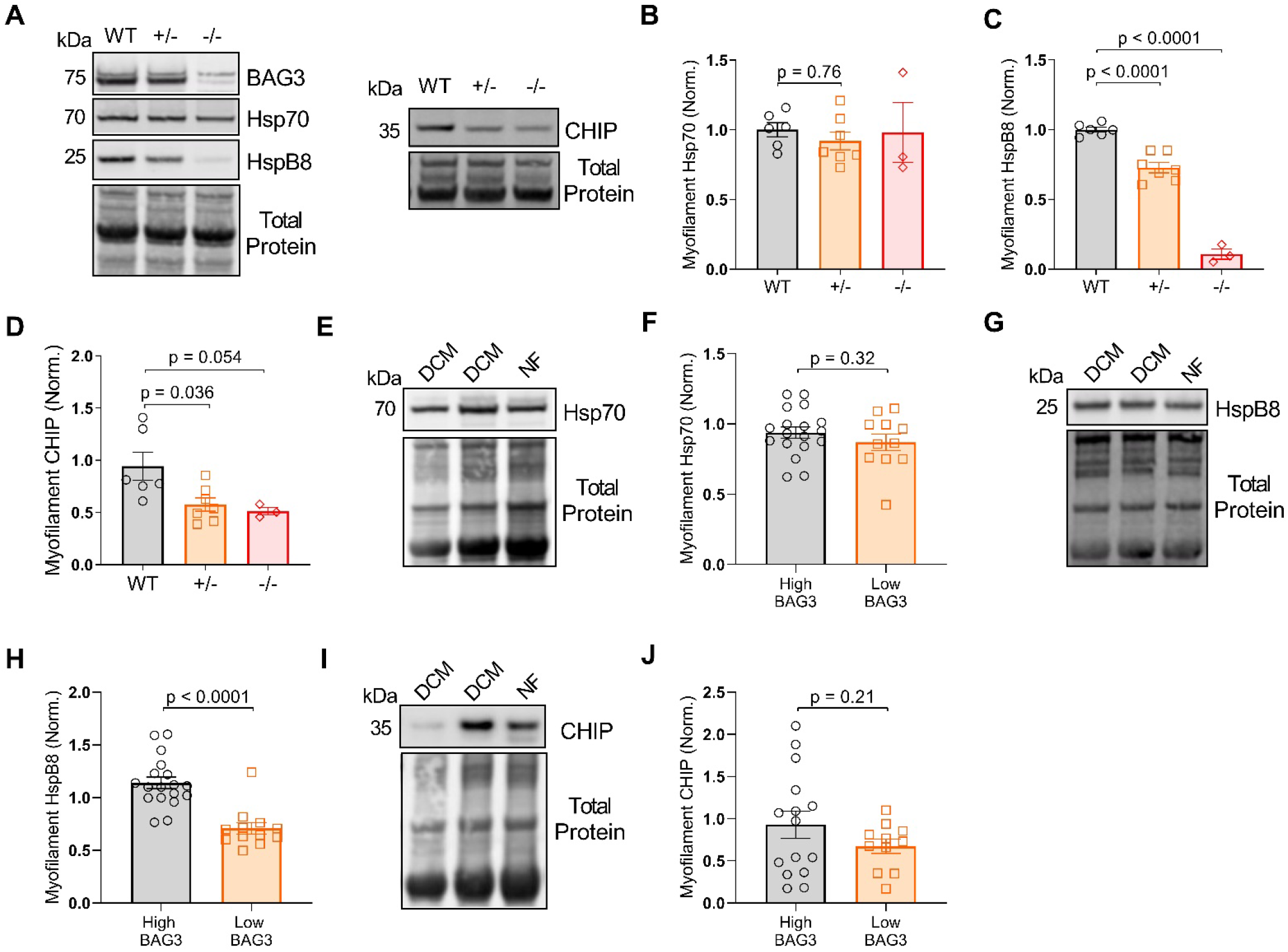
BAG3 is required for full assembly of CASA complex members in the myofilament fraction. **A.** Representative western blots for myofilament BAG3, Hsp70, HspB8, and CHIP in WT, BAG3^+/-^, and BAG3^-/-^ mice. **B-D.** Myofilament expression of Hsp70 (B), HspB8 (C), and CHIP (D) normalized to total protein loading control; n = 6 WT, 7 BAG3^+/-^, 3 BAG3^-/-^; one-way ANOVA, Tukey post-hoc. **E.** Representative western blot for myofilament Hsp70 in DCM and NF human samples. **F.** Myofilament Hsp70 expression in the human samples grouped by “high” or “low” BAG3 expression. **G-H.** Representative western blot for myofilament HspB8 in DCM and NF human samples (G) and grouped by “high” or “low” BAG3 expression (H). **I-J.** Representative western blot for myofilament CHIP in DCM and NF human samples (I) and grouped by “high” or “low” BAG3 expression (J). High ≥ 80% of mean NF expression, Low < 80% mean NF expression; 2-tailed *t*-test. All data are presented as mean ± SEM.

Since myofilament BAG3 expression decreases in human DCM, we next sought to determine whether HspB8 and CHIP display a similar relationship to BAG3 in humans as in the BAG3^+/-^/BAG3^-/-^ mice. As shown in Figure 2, the BAG3^+/-^ mice have ~80% of the wild-type myofilament BAG3, which we found to be a functionally significant decrease in expression. Therefore, we separated the human samples into two groups: those with “high” BAG3 expression (≥80% of the mean NF BAG3 expression) and those with “low” BAG3 expression (<80% of the mean NF BAG3 expression). As in the mice, Hsp70 did not display BAG3-dependent localization to the myofilament (**Figure 5E-F**). However, patients with lowest BAG3 expression had significantly reduced myofilament HspB8 levels (**Figure 5G-H**). Myofilament CHIP expression in the humans also trended toward a decrease in the low BAG3 group, though there was considerable variability (**Figure 5I-J**). Together, our data in the mouse and human tissue support that BAG3 is required for complete assembly of the CASA complex members at the myofilament, either through directly mediating HspB8/CHIP localization to the myofilament or serving to stabilize these proteins and prevent their degradation, which is supported for HspB8 by a previous study^24^.

### BAG3 gene therapy in heart failure restores F_max_ and reduces myofilament ubiquitination

Having established BAG3 is crucial for maintaining sarcomere function and identifying CASA as a potential mechanism, we next hypothesized that increasing BAG3 expression in heart failure through adenovirus gene therapy would rescue myofilament force-generating capacity through restored CASA. Eight-week-old mice received either a myocardial infarction produced by permanent coronary artery ligation (HF) or sham surgery (Sham). Eight weeks post-surgery, the mice were randomly assigned to receive a recombinant adeno-associated virus serotype 9 (AAV9) vector expressing the mouse *bag3* gene or control (AAV9/GFP) via retro-orbital injection. Four weeks after AAV injection the mice were euthanized, and the LV tissue was collected (**Figure 6A**). BAG3 overexpression in heart failure restored *in vivo* cardiac function to sham levels, which was reported previously for this cohort^44^.

**Figure 6.**
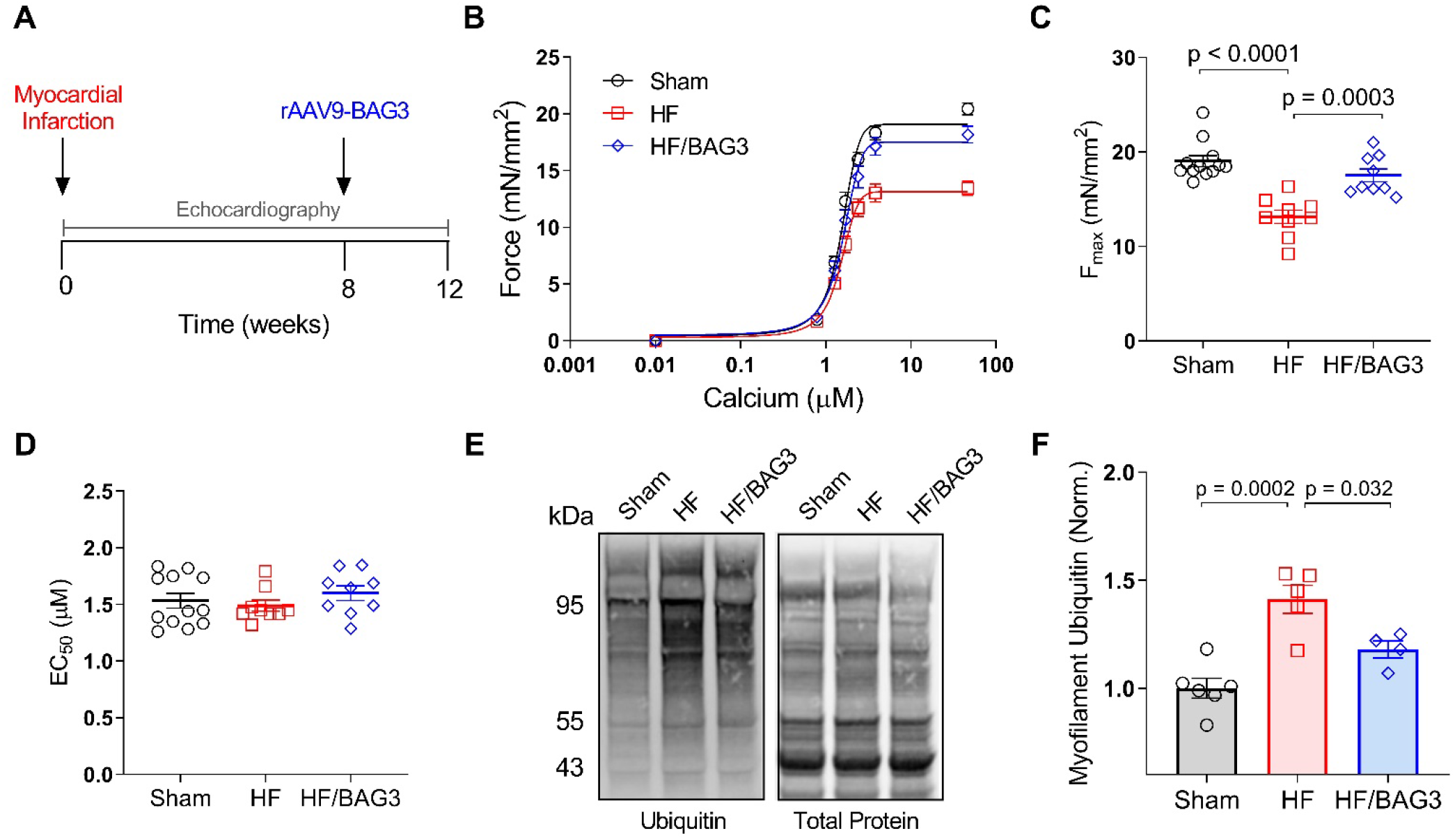
Increasing BAG3 expression in heart failure rescues myofilament function and restores sarcomere proteostasis. **A.** Experimental paradigm: 8-week-old mice received myocardial infarction by LAD ligation to induce progression into heart failure. Eight weeks post-MI, mice received either AAV9/GFP or AAV9/BAG3 by retro-orbital injection. Four weeks later mice were euthanized. **B.** Skinned myocyte force-calcium relationship from sham, HF, and HF/BAG3 cardiomyocytes. **C-D.** Summary data for myocyte F_max_ (C) and EC_50_ (D); n = 12 sham from 4 mice, 9 HF from 3 mice, and 9 HF/BAG3 from 3 mice. **E.** Representative western blot for myofilament ubiquitin in the sham, HF, and HF/BAG3 mice. **F.** Myofilament ubiquitin normalized to total protein loading control; n = 6 sham, 5 HF, 4 HF/BAG3. All data were analyzed via one-way ANOVA with Tukey post-hoc test for multiple comparisons. Data are presented as mean ± SEM.

We assessed myofilament function using force-calcium experiments on skinned myocytes from the infarct border zone. As in human DCM, cardiomyocytes from mice in the HF group displayed significantly reduced F_max_. However, in the HF mice that received AAV9/BAG3, F_max_ was fully restored to Sham levels (**Figure 6B-C**). Changes in myofilament calcium sensitivity were not observed, which may be attributed to the relatively early stage of disease progression we studied (**Figure 6D**). To determine the impact of BAG3 gene therapy on sarcomere proteostasis, we next assessed myofilament protein ubiquitination between the three groups by western blot. We found ubiquitin expression in the myofilament fraction increased significantly in the HF mice but was reduced with AAV9/BAG3 (**Figure 6E-F**). Together, these data support that increasing BAG3 expression in heart failure rescues myofilament force-generating capacity by restoring sarcomere protein turnover.

### BAG3 gene therapy in HF restores turnover of several sarcomere proteins

Because myofilament ubiquitination increased in the HF mice, we next tested whether this was due to a failure of the CASA complex to localize to the myofilament fraction. Using western blot, we found instead that Hsp70, HspB8, and CHIP levels were each increased at the myofilament in the HF mice (**Figure 7A-H**). These results indicate that targeting of the complex to the myofilament in response to proteotoxic stress is not hindered in the early progression to heart failure. Instead, the clearance of the CASA complex and its ubiquitinated clients appears to be impaired, leading to sarcomere proteotoxicity and decreased contractile function. The issue of CASA clearance was rectified by BAG3 gene therapy, which restored expression of all CASA members to Sham levels (**Figure 7A-H**). Notably, myofilament levels of the autophagic ubiquitin receptor P62/SQSTM1 also increased in the HF mice and were returned to baseline with BAG3 gene therapy (**Figure 7G**). P62 is an essential adaptor protein for autophagy that mediates the targeting of ubiquitinated proteins to the autophagosome for degradation^45^. Thus, its aggregation at the myofilament further suggests impaired autophagic clearance by CASA in heart failure that was rescued by AAV9/BAG3 gene therapy.

**Figure 7.**
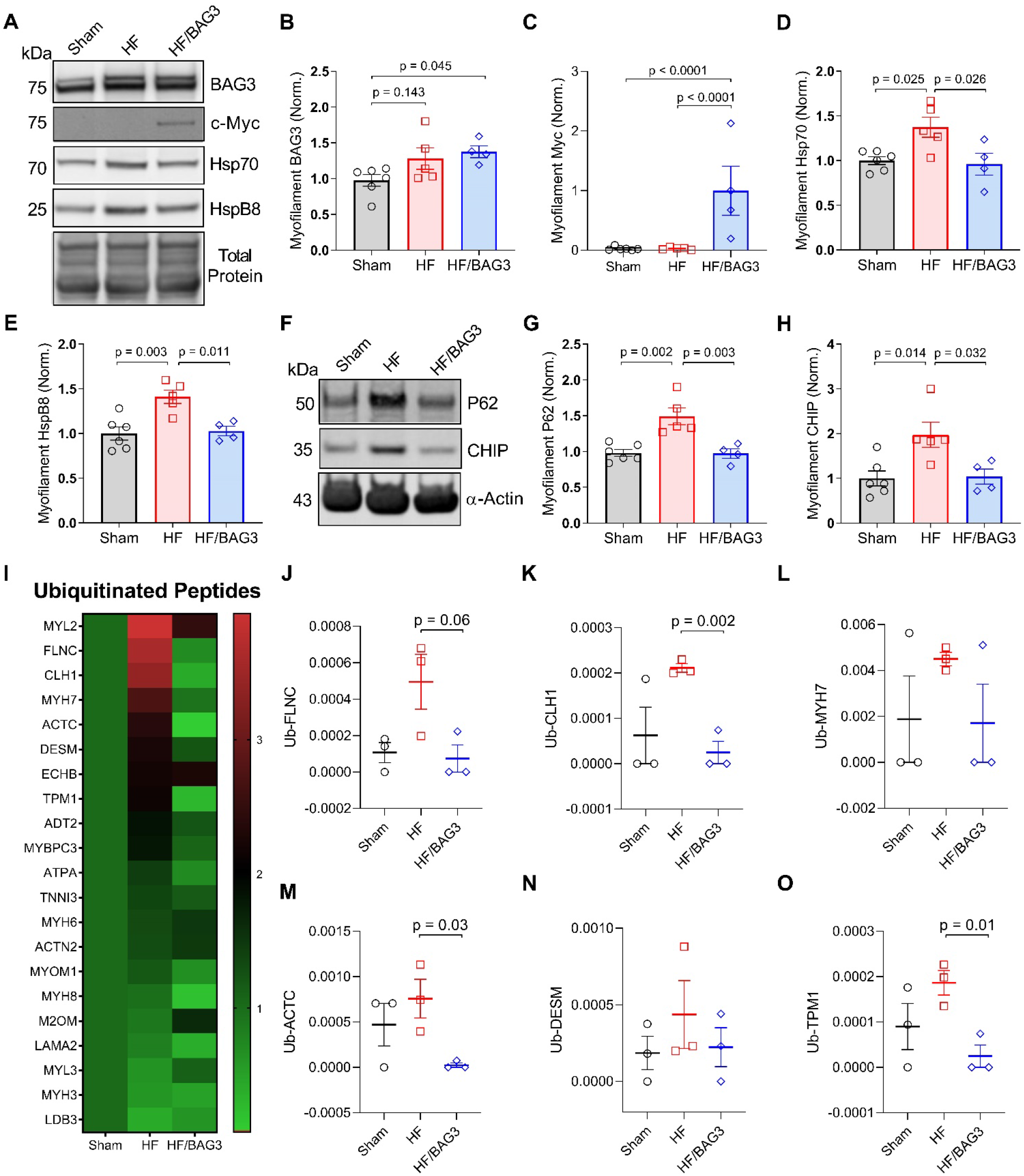
CASA clearance at the sarcomere stalls in heart failure but is restored by BAG3 gene therapy, which increases sarcomere protein turnover. **A-E.** Representative western blot for myofilament BAG3, *myc*-BAG3, Hsp70, and HspB8 (A) and normalization to total protein loading control (B-E) in the sham, HF, and HF/BAG3 mice. **F-H.** Representative western blot for myofilament P62 and CHIP (F) and normalized to total protein loading control (G-H); n = 6 sham, 5 HF, 4 HF/BAG3. **I.** Heat map of ubiquitinated proteins identified by LC-MS/MS; scale bar = fold change relative to Sham. **J-O.** Spectral count data for ubiquitinated peptides normalized to total peptide input for filamin C (J), clathrin heavy chain (K), β-MHC (L), actin (M), desmin (N), and tropomyosin (O); n = 3 samples/group. All data are presented as mean ± SEM and were analyzed by one-way ANOVA, Tukey post-hoc.

To specifically characterize how BAG3 gene therapy improved sarcomere function, we next sought to identify which sarcomere proteins are substrates of BAG3-dependent degradation. Our initial mass spectrometry assessment of BAG3-associated myofilament proteins identified many binding partners, of which some may be CASA clients (**Supplemental Table 1**). However, this approach was not designed to determine which of the BAG3-associated proteins were ubiquitinated, and thus possible substrates. Therefore, in order to identify the sarcomeric CASA clients, we performed mass spectrometry on ubiquitin-enriched peptides from the Sham, HF, and HF/BAG3 mice, similar to the experiment we did on human samples in Figure 1, anticipating that CASA clients would be increased in HF and reduced to baseline in the HF/BAG3 group. Our LC-MS/MS analysis identified several ubiquitinated sarcomere proteins that were reduced with BAG3 gene therapy (**Figure 7I**). Foremost among these were the previously proposed CASA client in skeletal muscle, filamin C (FLNC), along with the novel clients sarcomeric α-actin (ACTC), tropomyosin (TPM1), and clathrin heavy chain 1 (CLH1), an important factor for sarcomere organization and autophagosome formation^46–48^. Other potential clients that did not reach significance due to high variability of ubiquitinated peptides identified include β-MHC (MYH7) and desmin (DESM), which tended to increase in heart failure and were reduced in the HF/BAG3 group (**Figure 7J-O**). Taken together, our results identify sarcomeric CASA is impaired in heart failure, however, increasing BAG3 expression via AAV9/BAG3 restores CASA clearance of a subset of ubiquitinated sarcomeric proteins.

## DISCUSSION

On the cellular level, decreased cardiac contractile strength in heart failure may be attributed to structural and functional changes at the sarcomere, the molecular unit of contraction in striated muscle^3^. However, the mechanisms underlying decreased sarcomere tension generation in heart failure are incompletely understood, especially for the long-observed depression in F_max_. In this study, we sought to determine if inadequate sarcomeric protein turnover in heart failure could serve as an explanation for functional impairment and, if so, to identify the key players involved. First, we show sarcomere protein turnover is impaired in human DCM samples with reduced myofilament force-generating capacity (F_max_), where elevated ubiquitinated (and thus presumably misfolded/dysfunctional) proteins remained integrated into the sarcomere structure. We found myofilament localization of the co-chaperone BAG3, which decreases in DCM, is predictive of the extent of myocyte F_max_ decrease in humans and that BAG3 haploinsufficiency in a mouse model reduced F_max_ and increased myofilament protein ubiquitination. The relationship between impaired BAG3-mediated protein degradation and decreased sarcomere function was substantiated using a transgenic mouse model with the P209L BAG3 mutation, which impairs ubiquitinated client release to autophagy. The P209L mice displayed a reduction in F_max_ and increased myofilament ubiquitination. We identify BAG3-mediated chaperone-assisted selective autophagy (CASA) as the mechanism of BAG3-mediated sarcomere turnover and show CASA is impaired in the progression to heart failure. However, CASA clearance in heart failure was restored by BAG3 gene therapy, which rescued turnover of several sarcomere proteins and increased F_max_. This study reveals BAG3-mediated sarcomere protein turnover, which is disrupted in heart failure, is fundamental for maintaining optimal sarcomere contractile function.

Disrupted proteostasis has long been associated with sarcomere structural abnormalities and heart failure^12,49–51^. However, mechanisms of sarcomere protein turnover and the myofilament functional significance of individual molecular chaperones involved in sarcomere protein quality control are not known. BAG3 is key to protein quality control in many different cell types and is involved in two separate autophagy pathways, chaperone mediated autophagy and selective macroautophagy^26,28,52^. Interestingly, BAG3 had previously been shown to localize to the sarcomere Z-disc, positioning it as a potential mediator of sarcomere protein turnover through autophagy^38^. However, relatively little was known regarding BAG3’s role at the cardiac sarcomere. Mechanistic studies in neonatal myocytes found BAG3 was required for sarcomere structural organization through stabilization of the actin-capping protein CapZ^21^. However, neonatal myocytes are structurally and functionally distinct from the adult cardiomyocyte and are not under any mechanical load, making inferences from this study to the adult difficult. Indeed, it is apparent that BAG3’s role in adult cardiomyocytes differs from the neonatal as BAG3 was found dispensable for sarcomere structural maintenance in utero, but loss of BAG3 caused rapid sarcomere disintegration once the heart was under mechanical load^23^. Moreover, recent studies in the adult heart failed to identify CapZ as a BAG3-interacting protein, suggesting the stabilization of CapZ by BAG3 is specific to sarcomere assembly^22^. Our mass spectrometry analysis of the myofilament BAG3 interactome also failed to identify CapZ as a binding partner, further suggesting this association is specific to development.

An important advance in our understanding of BAG3’s role in the adult heart came from a study by Fang *et al*. who showed that cardiomyocyte-restricted BAG3 KO caused a DCM phenotype in mice, was associated with reduced stability of the small heat shock proteins (HspBs), and increased toxic protein aggregation^24^. The authors also showed that a DCM-associated BAG3 mutation (E455K) impaired BAG3 binding to Hsp70 and was associated with reduced stability of HspBs and protein aggregation. Together, these findings strongly indicate BAG3 is required for protein turnover in the adult heart. However, while this study identified prominent protein aggregation with decreased/dysfunctional BAG3, it did not assess whether misfolded proteins remained incorporated into the sarcomere. In the present study, we found that sarcomere-specific expression of BAG3 decreased in human DCM and levels were particularly low in samples with the weakest myofilament force-generating capacity. We therefore hypothesized that impaired BAG3-mediated turnover of sarcomere proteins might contribute to decreased sarcomere contractile function. Using cardiomyocyte-restricted BAG3 haploinsufficiency, we found elevated sarcomere protein ubiquitination, suggesting inadequate turnover, and reduced myofilament force-generating capacity. Importantly, these ubiquitinated proteins were not members of protein aggregates, but were still embedded within the sarcomere complex and did not cause overt structural disarray, thus offering a compelling and novel explanation for the observed functional deficit.

To further clarify the functional importance of BAG3-mediated protein turnover for the sarcomere, we used a transgenic mouse model expressing the human P209L mutation. The P209L BAG3 mutation is highly penetrant and is associated with restrictive cardiomyopathy, myofibrillar myopathy, and elevated toxic protein aggregation, which present in early adolescence^20,36^. Recent work in cell culture systems identified the P209L mutation impairs release of ubiquitinated substrates to the macroautophagy system^40,53^. We chose transgenic expression of the human mutant BAG3 as a recent study of the analogous mouse mutation (P215L) showed no cardiac phenotype, which was attributed to the pathogenicity of this variant being specific to the human BAG3 isoform^54^. Previous work with the human P209L transgenic model found mice developed restrictive cardiomyopathy by 8 months, mimicking the human phenotype^41^. In the present study, we found cardiac expression of the P209L BAG3 mutant increased myofilament ubiquitin levels and caused a significant reduction in F_max_ by 8 months of age. Instead of mediating sarcomere stability as in the neonatal myocyte, our data identify a converse role for BAG3 in the adult myocyte, where BAG3 is required for sarcomere protein turnover.

Studies in skeletal muscle previously identified BAG3 mediates chaperone-assisted selective autophagy (CASA), a macroautophagy pathway involving Hsp70 and HspB8^26,27^. CASA is a ubiquitin-dependent autophagy pathway whereby ubiquitinated proteins are delivered to the autophagosome and degraded by lysosomal hydrolases upon autophagosome-lysosome fusion^28^. In skeletal muscle, CASA mediates the degradation of the actin cross-linking protein filamin C when it is denatured due to mechanical strain^26,27^. However, CASA had never been previously described in the heart. We identified the BAG3/Hsp70/HspB8 complex assembles at the sarcomere Z-disc in a BAG3- and proteotoxic stress-dependent manner. We further show the CASA members, the E3 ubiquitin ligase for CASA substrates (CHIP), and the autophagic ubiquitin receptor P62, aggregate at the myofilament in the early progression into heart failure, suggesting impaired autophagic clearance^55^. However, when we increased BAG3 expression using AAV-mediated gene therapy, myofilament expression of these proteins returned to normal levels. Using ubiquitinated peptide enrichment for mass spectrometry, we identified filamin C, sarcomeric α-actin, clathrin heavy chain, and tropomyosin as myofilament CASA clients which were reduced by AAV9-BAG3 treatment in heart failure.

The identification of tropomyosin as a CASA client is of interest, as it directly modulates sarcomere contraction and was also among the top proteins with increased ubiquitination in human DCM. Tropomyosin regulates the cross-bridge cycle by forming dynamic interactions along the thin filament, which at low calcium concentrations prohibit the interaction of myosin heads with actin^56^. Under normal circumstances, increases in calcium cause the tropomyosin chain to shift, exposing myosin binding sites on actin, and facilitating sarcomere contraction. The increased ubiquitination of tropomyosin that we identified in heart failure indicates tropomyosin has been marked for degradation, however, it is not adequately removed from the sarcomere. We expect that such an accumulation of misfolded/dysfunctional tropomyosin remaining integrated into the sarcomere in heart failure causes contractile impairment by prohibiting proper cross-bridge regulation. The link between tropomyosin and F_max_ is supported by numerous earlier biophysical studies that indicate tropomyosin is fundamental for cross-bridge force generation^57–62^. Thus, inadequate turnover of tropomyosin serves as a possible explanation for decreased sarcomere force generating capacity in heart failure, each of which were ameliorated by increasing BAG3 expression in the present study.

Considered as a whole, we identify inadequate sarcomere protein turnover as a mechanism of sarcomere functional decline in heart failure. Moreover, our findings show decreased BAG3-mediated sarcomere protein degradation is a primary contributor to this observed decrease in myofilament contractile function. We confirm the importance of BAG3-mediated filamin C degradation (previously identified in skeletal muscle) in cardiomyocytes and identify three new clients: actin, tropomyosin, and clathrin heavy chain. These proteins are fundamental for myofilament contraction and sarcomere organization, and we thus propose that their impaired BAG3-mediated turnover in heart failure causes the sarcomere functional deficit observed. Notably, the four BAG3/CASA clients are all thin filament or thin filament-associated proteins. Thus, BAG3-mediated degradation of the sarcomere thin filament emerges as a novel mechanism of sarcomere functional maintenance in the adult cardiomyocyte. Further work is needed to determine whether BAG3 mediates degradation of β-MHC and desmin, which is suggested from our data. This is the first study to indicate impaired protein quality control at the sarcomere in heart failure has a direct effect on myofilament function. Furthermore, our results show BAG3 is required for maintaining sarcomere proteostasis and optimal tension generation in cardiomyocytes and demonstrate the potential therapeutic benefits of BAG3 gene therapy in heart failure include improving myofilament function by restoring sarcomeric protein turnover.

## METHODS

### Human Heart Tissue Procurement

Human left ventricular tissue was obtained from the Loyola Cardiovascular Research Institute Biorepository and Cleveland Clinic Biorepository. Tissue from failing human hearts with non-ischemic idiopathic dilated cardiomyopathy and left ventricular dysfunction was collected either at the time of heart explant or at the time of LVAD implantation. Informed consent was obtained prior to tissue collection. Myocardial tissue from the non-failing donors with no history of coronary artery disease or heart failure (EF%, 57.5 ± 3.2) was collected post-mortem with other organs. All tissue was flash frozen in liquid nitrogen. The patient age and sex distribution were not significantly different between the non-failing (53.3 ± 8.9 years, 44.4% female) and failing hearts (55.5 ± 11.5 years, 38.1% female).

### Mouse Models

#### BAG3 Overexpression Model

The generation of mice with LV dysfunction secondary to a left coronary artery ligation was described previously^44^. In brief, eight-week old C57/Bl6 mice were randomized to undergo a left descending coronary artery (LAD) ligation or sham surgery^63^. Because only half of the mice in which the artery was ligated were alive at one-week post-surgery, mice were not randomized to treatment group until one-week post-surgery. Eight weeks post-MI, the now randomized sham-operated mice and post-infarction mice received either gene therapy with a recombinant adeno-associated virus-9 expressing BAG3 (rAAV9-BAG3) or control (AAV9-GFP) by retro-orbital injection. Four weeks later mice were sacrificed, tissue was removed, and flash frozen in liquid nitrogen.

#### BAG3 P209L Model

The generation of the P209L BAG3 mouse strain was described previously^41^. Transgene positive founders were bred into wild-type C57/Bl6 mice. P209L positive offspring were identified through PCR of isolated tail DNA obtained during weaning using primers for the human BAG3 transgene. Wild-type and transgene-positive offspring were pair-housed and aged to 3 or 8 months, at which time they were euthanized, and myocardial tissue was collected.

#### BAG3 +/- & -/- Model

Generation of mice homozygous (-/-) or heterozygous (+/-) for the murine version of the constitutive (c) deletion of *BAG3* (*cBAG3^-/-^* or *cBAG3^+/-^*) has been described in detail elsewhere^39^. Mice lacking a single allele of wild type *bag3* developed early LV dysfunction by 8 to 10 weeks of age. Mice in which both alleles were deleted developed LV dysfunction commensurate with that of the heterozygous mice; however, the BAG3^-/-^ died by 12 to 14 weeks of age with severe LV dysfunction. Hemodynamic data for the *BAG3^+/-^* was reported previously^39^. Hemodynamic data for the BAG3^-/-^ was obtained separately. The mice used for data collection in this study were 6 weeks of age, thus preceding the development of detectable *in vivo* LV dysfunction.

### Skinned Myocyte Force-Ca^2+^ Experiments

Skinned cardiomyocytes were prepared as described previously^64^. Briefly, frozen left-ventricular tissue was placed in Isolation solution (**Supplemental Table 2**) containing protease and phosphatase inhibitors (1:100 ratio, Fisher Scientific), 10 mM 1,4-dithiothreitol (DTT, Millipore Sigma), and 0.3% (v/v) Triton X-100 (Millipore Sigma). The tissue was mechanically homogenized in three one-second bursts at 7,000 RPM and left on ice for 20 minutes to permeabilize. Next, the myocyte solution was pelleted by centrifugation at 120*g* for 2 minutes and resuspended in Isolation solution without triton. To assess function, myocytes were first attached with UV-curing glue (NOA 61, Thorlabs) to two pins, one attached to a force transducer (AE-801, Kronex) and the other to a high-speed piezo length controller (Thorlabs). Next, the myocyte was perfused with a maximal calcium Activating solution to elicit contraction and then perfusion was switched to 100% Relax solution (**Supplemental Table 2**) to facilitate relaxation. This process was continued with perfusion of five additional submaximal calcium solutions containing Activating solution mixed with Relax solution (**Figure S4**). All measurements were conducted at room temperature and performed at a sarcomere length of 2.1 μm as measured by Fast Fourier Transform. Experimenters were blinded to the experimental groups during the collection of these data.

### Neonatal Rat Ventricular Myocyte Studies

Neonatal rat hearts were isolated from 0-1 day old Sprague-Dawley rat pups, placed in ice cold Krebs-Henseleit Buffer (KHB)^65^, and rinsed twice. Next, the atria were removed, and the isolated ventricles were digested in 37 °C KHB supplemented with 1g/100mL collagenase type-II (Worthington) and 600 μl of 2.5% Trypsin/100mL (Gibco) with intermittent stirring. The buffer solution was then moved to a tube containing Dulbecco’s Modified Eagle’s Medium (DMEM, Gibco) supplemented with 10% fetal bovine serum (FBS, Millipore Sigma) to stop the digestion reaction. This process was repeated four times or until the myocardial tissue was fully digested and turned white. Next, the digested ventricular myocyte solution was centrifuged at 140*g* for 10 minutes and the pelleted cells were resuspended in DMEM with 10% FBS, ITSx, and penicillin/streptomycin, plated, and incubated at 37 °C in a 5% CO_2_ incubator for 90 minutes to allow the fibroblasts to stick. After incubation the plates were softly tapped to dislodge cardiomyocytes, the supernatant containing the cardiomyocytes was collected and cultured at 1 million cells per plate on 60 x 15 mm culture dishes coated with 0.1% gelatin and incubated at 37 °C/5% CO_2_. After 48 hours, the media was replaced with fresh media containing 2 μM MG132 (MedChem Express) dissolved in DMSO to induce proteotoxic stress by inhibiting the proteasome, or equal volume of DMSO (Millipore Sigma) vehicle control. After 24 hours of treatment the cells were collected and enriched for the myofilament-specific fraction.

### Myofilament Enrichment

Frozen left ventricular tissue was placed in 1 mL of a standard rigor buffer (SRB)^66^ containing protease and phosphatase inhibitors (1:100 ratio) and 0.5% Triton X-100. The tissue homogenized with a mechanical homogenizer and left on ice for 20 minutes. For NRVMs, the cells were suspended in SRB with Triton and left to permeabilize on ice for 20 minutes without homogenization. Myofilaments were pelleted by centrifugation at 1,800*g* for 2 minutes and the supernatant containing soluble proteins was removed. The myofilament pellet was washed twice in SRB without Triton, resuspended in 9M Urea (Millipore Sigma), and sonicated. Following sonication, the solution was centrifuged at 12,000*g* for 10 minutes and the supernatant containing the now solubilized myofilament proteins was collected.

### Immunoblotting

After isolating the myofilament fraction, protein concentration was determined using a BCA assay (Thermo Fisher). Samples were prepared in SDS Tris-Glycine Buffer (Life Technologies) supplemented with Bolt Reducing Buffer (Fisher Scientific), heated for 10 minutes at 95 °C, and run on 4-12% gradient Tris-glycine gels (Invitrogen). The proteins were transferred onto a nitrocellulose membrane (Thermo Scientific). Following transfer, membranes were incubated with Revert Total Protein Stain (LI-COR Biosciences) for five minutes to assess equal loading and then blocked in a 1:1 ratio Intercept Blocking Buffer (LI-COR Biosciences) to Tris-Buffered Saline (TBS) solution supplemented with 0.1% tween for 1 hour at room temperature. The following primary antibodies and dilutions were used and incubated in blocking buffer overnight at 4 °C with gentle rocking: BAG3 (Proteintech, 10599-1AP, 1:5,000), BAG3 (Santa Cruz Biotechnology, sc-135437, 1:500), c-Myc (Cell Signaling Technology, 71D10, 1:1000), HspB8 (Proteintech, 15287-1AP, 1:3,000), HspB8 (R&D Systems, MAB4987, 1:800), Hsp70 (Proteintech, 10995-1AP, 1:3,000), Ubiquitin (Cytoskeleton Inc., AUB01, 1:400), CHIP/STUB1 (Santa Cruz Biotechnology, sc-133083, 1:100), P62 (Proteintech, 18420-1AP, 1:3,000). Blots were washed in TBS-T and then incubated with IRDye secondary antibodies (LI-COR) in blocking solution for 1 hour with agitation at room temperature. Blots were analyzed using the LI-COR Image Studio software.

### Co-Immunoprecipitation

Dynabeads Protein G (Invitrogen) were incubated with 5 μg of antibody in 200 μl TBS with 0.1% Tween (TBS-T) for 20 minutes at room temperature with rotation. The immunocomplex was washed three times with 100 mM sodium borate (pH 9) and then antibody was crosslinked with 20 mM dimethyl pimelimidate dihydrochloride (DMP) in 100 mM sodium borate for 30 minutes at room temperature. After crosslinking, the complex was blocked with 200 mM ethanolamine (pH 8) for two hours. After blocking, the immunocomplex was washed three times with IAP buffer (50 mM MOPS (pH 7.2), 10 mM sodium phosphate, 50 mM NaCl) and then resuspended in 200 μl of 2 μg/μl myofilament-enriched protein lysates. After incubation for 10 minutes at room temperature with rotation, the immunocomplex was washed seven times with TBS-T on a magnetic rack. Immunoprecipitated proteins were eluted by suspending the beads in a 0.15% TFA solution for 5 minutes at room temperature. The eluted proteins were dried by vacuum centrifugation and then prepared for mass spectrometry or analyzed by western blot.

### Mass Spectrometry

#### In-solution digestion

Following immunoprecipitation for BAG3 the co-immunoprecipitated protein was reconstituted in 50 mM Tris-HCl (pH 8) and reduced with 5 mM DTT for 45 minutes. Following reduction, the proteins were alkylated by incubation in 10 mM iodoacetamide for 30 minutes. The proteins were next digested with 5 ug Trypsin/LysC protease for 18 hours at 37 °C/660RPM. The reaction was stopped by adding TFA at 0.2% final volume to bring the pH under 3. Samples were then dried by vacuum centrifugation, reconstituted in 300 μl of 0.1% TFA, and separated into 12 fractions by basic pH reverse phase fractionation. The fractions were concatenated, dried, reconstituted in buffer A (3% acetonitrile, 0.1% formic acid) and analyzed by liquid chromatography tandem mass spectrometry (LC-MS/MS).

#### Ubiquitinated Myofilament Peptide Enrichment

This procedure was adopted from Udeshi *et al*.^67^ with minor modifications. Approximately 500 μg of mouse myofilament protein, or 2 mg of human myofilament protein, were incubated in 5 mM final concentration DTT at room temperature for 45 minutes to reduce disulfide bonds, followed by incubation with 10 mM IAA for 30 minutes to carbamidomethylate cysteine residues. The protein was then diluted 1:4.5 with 50 mM Tris-HCl (pH 9) to reduce the urea concentration and digested with Trypsin/LysC protease mix (1:50 wt/wt) for 18 hours at 37 °C (pH 8-8.5) on a shaking incubator at 600 RPM. The peptides were next desalted with a 100-mg tC18 SepPak cartridge (Waters), dried by vacuum centrifugation, and stored at −80 °C until next use. Dried peptides were reconstituted in 0.1% TFA and separated into 12 fractions via high pH reverse phase fractionation (Pierce). At this stage a small amount of sample was set aside to be run as a total peptides control on LC-MS/MS later. The collected fractions were pooled by concatenation to maximize LC gradient utilization and then dried by vacuum centrifugation until next use.

Ubiquitinated peptides were purified using immunoaffinity purification with the diglycine-lysine ubiquitin remnant motif antibody (Cell Signaling Technology, PTMScan) cross-linked to Protein A agarose beads. Each pooled fraction was reconstituted in ice-cold IAP buffer (50 mM MOPS pH 7.2, 10 mM NaPO_4_ dibasic, 50 mM NaCl) and incubated with 30 μg of antibody for 1 hour at 4 °C with gentle rotation. After washing, the purified peptides were eluted by incubation with 0.15% TFA and desalted using the GL-Tip SDB 200 μl C18 tips (GL Biosciences). The collected samples were then dried by vacuum centrifugation and stored at −80 °C until ready for LC-MS/MS.

Tryptic peptides were reconstituted in 20 μl of buffer A (3% ACN, 0.1% FA) and 5 μl were subjected to high pressure liquid chromatography (HPLC) in a 25 cm PepMap RSLC C18 column (Thermofisher) coupled to tandem mass spectrometry (MS/MS) on an LTQ Orbitrap XL mass spectrometer. Upon completion, raw data files were imported into the Peaks Bioinformatics program and acquired masses were searched against the *Homo sapiens* or *Mus musculus* database. The precursor ion mass tolerance was set at 20 ppm and the MS/MS was set at 0.5 Da. A maximum of three missed cleavages were allowed. The fixed modification was carbamidomethylated cysteine (+57.02 Da) and the variable modification was ubiquitinated lysine (+114.04 Da). For comparison between groups, ubiquitinated peptides were normalized to the corresponding total peptides control for each sample set aside earlier.

### Immunofluorescence microscopy

Frozen left ventricular tissue was placed in Isolation solution (Supplemental Table 2) containing 0.3% Triton X-100, homogenized in 3-4 one-second bursts at 7,000 RPM with a mechanical homogenizer, and left for 20 minutes on ice to permeabilize. Next, the cells were pelleted by centrifugation at 120*g* for 2 minutes, resuspended in Isolation solution without triton, and seeded on Poly-L-Lysine-coated (Millipore Sigma) chamber slides (Thermo Scientific). Once settled, the cells were fixed with ice-cold methanol for 1 minute and 4% paraformaldehyde for 3 minutes. For NRVMs, freshly isolated myocytes were plated on nanopatterned coverslips (Nanosurface Cultureware) to promote adult-like myocyte morphology. Forty-eight hours after plating, the myocytes were treated with 2 μM MG132 or equal volume of DMSO. After 24 hours, the myocytes were fixed with ice cold methanol for 1 minute, followed by 4% paraformaldehyde for 3 minutes.

Fixed myocytes were incubated in 0.5% Triton solution for 20 minutes, followed by 0.1% Triton for 30 minutes to permeabilize the sarcolemma. Following permeabilization, the cells were incubated in 0.1M glycine antigen retrieval solution for 30 minutes, washed three times with PBS, and incubated for one hour in BSA blocking buffer at room temperature. Next, primary antibodies were added in blocking solution and incubated for 12-14 hours at 4 °C. Primary antibodies and dilutions used: BAG3 (Proteintech, 10599-1AP, 1:300), HspB8 (Proteintech, 15287-1AP, 1:300), Hsp70 (Proteintech, 10995-1AP, 1:300), CHIP/STUB1 (Proteintech, 55430-1-AP, 1:300), P62 (Proteintech, 18420-1AP, 1:250), α-actinin (Millipore Sigma, A7811, 1:250), oligomer A11 (Invitrogen, AHB0052, 1:100), Ubiquitin (Cytoskeleton Inc., AUB01, 1:40). After washing four times with PBS, the cells were incubated for one hour with the appropriate secondary antibody conjugated to Alexa-Fluor 488 or Alexa-Fluor 568 (Abcam, 1:1,000). Slides were mounted with Vectashield (Vector Laboratories), sealed with coverslips, and imaged at 63x magnification on a Zeiss LSM 880 super-resolution confocal microscope. Images were acquired under constant laser intensity and photomultiplier gain settings.

### Statistics

Data are presented as mean ± SEM and were analyzed using one-way ANOVA. When a significant variance was identified, Tukey’s post-hoc test for multiple comparisons was performed. Comparisons of two groups were made by 2-tailed Student’s *t* test for unpaired samples (GraphPad Prism 8).

### Study Approval

The experimental procedures used in the mouse protocols were approved by the Temple University and Hines VA IACUC. Temple University and Hines VA are accredited institutions recognized by the Association for Assessment and Accreditation of Laboratory Animal Care (AAALAC). Human samples used for this study were obtained from the tissue biorepositories at the Cleveland Clinic and the Cardiovascular Research Institute at Loyola University Chicago. Informed consent was obtained prior to tissue collection, which was performed with permission from the institutional review boards of Loyola University Chicago and the Cleveland Clinic.

## AUTHOR CONTRIBUTIONS

T.G.M and J.A.K designed the experiments. T.G.M performed the experiments. V.D.M, P.D, S.D, and E.P maintained mouse colonies and performed tissue collection. C.S.M provided human samples. M.S.W provided the BAG3_P209L_ mice. A.M.F provided tissue from the AAV9/BAG3 and BAG3^+/-^/BAG3^-/-^ mouse models. J.A.K provided scientific input from the conception of the project idea through its completion, and substantially contributed to the data presentation. T.G.M and J.A.K wrote the manuscript with input from all authors.

## ACKNOWLEDGEMENTS

We thank Dr. J.R Beach from Loyola University Chicago for providing access to the Zeiss LSM 880 in his laboratory, and Peter Caron from Loyola University Chicago for manufacturing custom pieces for our biophysical rigs.

