## Supplemental Methods, Figures, and Tables for "Cardiomyocyte Contractile Impairment in Heart Failure Results from Reduced BAG3-mediated Sarcomeric Protein Turnover"

**Supplemental Table 1. Myofilament BAG3 interactome as identified by LC-MS/MS from myofilament BAG3 immunoprecipitation.**

| Accession | Coverage (%) | #Peptides | #Unique | Avg. Mass |
|---|---|---|---|---|
| P12883\|MYH7_HUMAN | 30 | 61 | 18 | 223095 |
| P13533\|MYH6_HUMAN | 20 | 40 | 1 | 223733 |
| O95817\|BAG3_HUMAN | 68 | 37 | 37 | 61595 |
| P68032\|ACTC_HUMAN | 58 | 25 | 1 | 42019 |
| P17661\|DESM_HUMAN | 47 | 24 | 17 | 53536 |
| P12882\|MYH1_HUMAN | 10 | 22 | 0 | 223143 |
| P63261\|ACTG_HUMAN | 34 | 13 | 1 | 41793 |
| P35609\|ACTN2_HUMAN | 15 | 12 | 11 | 103854 |
| Q14896\|MYPC3_HUMAN | 10 | 11 | 11 | 140762 |
| Q9UJY1\|HSPB8_HUMAN | 44 | 10 | 10 | 21604 |
| A7E2Y1\|MYH7B_HUMAN | 4 | 9 | 2 | 221386 |
| P12235\|ADT1_HUMAN | 19 | 7 | 2 | 33065 |
| Q562R1\|ACTBL_HUMAN | 17 | 7 | 2 | 42003 |
| Q6S8J3\|POTEE_HUMAN | 6 | 7 | 0 | 121363 |
| P25705\|ATPA_HUMAN | 11 | 6 | 6 | 59751 |
| P08590\|MYL3_HUMAN | 39 | 6 | 5 | 21932 |
| P61353\|RL27_HUMAN | 28 | 5 | 5 | 15798 |
| P12236\|ADT3_HUMAN | 12 | 5 | 0 | 32866 |
| P0CG38\|POTEI_HUMAN | 4 | 5 | 0 | 121282 |
| Q9BYX7\|ACTBM_HUMAN | 12 | 5 | 0 | 42016 |
| P08670\|VIME_HUMAN | 8 | 5 | 1 | 53652 |
| P20674\|COX5A_HUMAN | 24 | 4 | 3 | 16762 |
| P63316\|TNNC1_HUMAN | 32 | 4 | 4 | 18402 |
| P14136\|GFAP_HUMAN | 5 | 3 | 0 | 49880 |
| P0DMV8\|HS71A_HUMAN | 5 | 3 | 0 | 70052 |
| Q9H6N6\|MYH16_HUMAN | 2 | 3 | 0 | 128290 |
| P09493\|TPM1_HUMAN | 12 | 3 | 1 | 32709 |
| Q9Y2D5\|AKAP2_HUMAN | 2 | 2 | 0 | 94661 |
| A6NCL7\|AN33B_HUMAN | 3 | 2 | 1 | 53975 |
| O75964\|ATP5L_HUMAN | 23 | 2 | 2 | 11428 |
| P31930\|QCR1_HUMAN | 5 | 2 | 2 | 52646 |
| P13073\|COX41_HUMAN | 12 | 2 | 2 | 19577 |
| Q8TD57\|DYH3_HUMAN | 0 | 2 | 1 | 470774 |
| P17066\|HSP76_HUMAN | 4 | 2 | 0 | 71028 |
| Q9NPC6\|MYOZ2_HUMAN | 9 | 2 | 2 | 29898 |
| Q13423\|NNTM_HUMAN | 2 | 2 | 1 | 113895 |
| Q9UHQ9\|NB5R1_HUMAN | 6 | 2 | 2 | 34095 |
| Q00325\|MPCP_HUMAN | 7 | 2 | 2 | 40095 |
| P48741\|HSP77_HUMAN | 7 | 2 | 0 | 40244 |
| Q15643\|TRIPB_HUMAN | 1 | 2 | 0 | 227584 |
| B2RTY4\|MYO9A_HUMAN | 1 | 2 | 1 | 292704 |

**Supplemental Table 2. Chemical composition of the Activating, Relaxing, and Isolation solutions used for skinned myocyte preparation and functional assessment.**

| Activating | Final Conc. (mM) | Relaxing | Final Conc. (mM) | Isolation | Final Conc. (mM) |
|---|---|---|---|---|---|
| $Ca^{2+}$-EGTA | 10 | EGTA | 10 | EGTA | 2 |
| Potassium Propionate | 28.1 | Potassium Propionate | 47.6 | Potassium Hydroxide | 8.9 |
| BES | 100 | BES | 100 | Imidazole | 10 |
| $MgCl_2$ | 6.2 | $MgCl_2$ | 6.5 | $MgCl_2$ | 7.1 |
| ATP | 6.3 | ATP | 6.2 | ATP | 5.8 |
| Creatine Phosphate | 10 | Creatine Phosphate | 10 | Potassium Chloride | 108 |

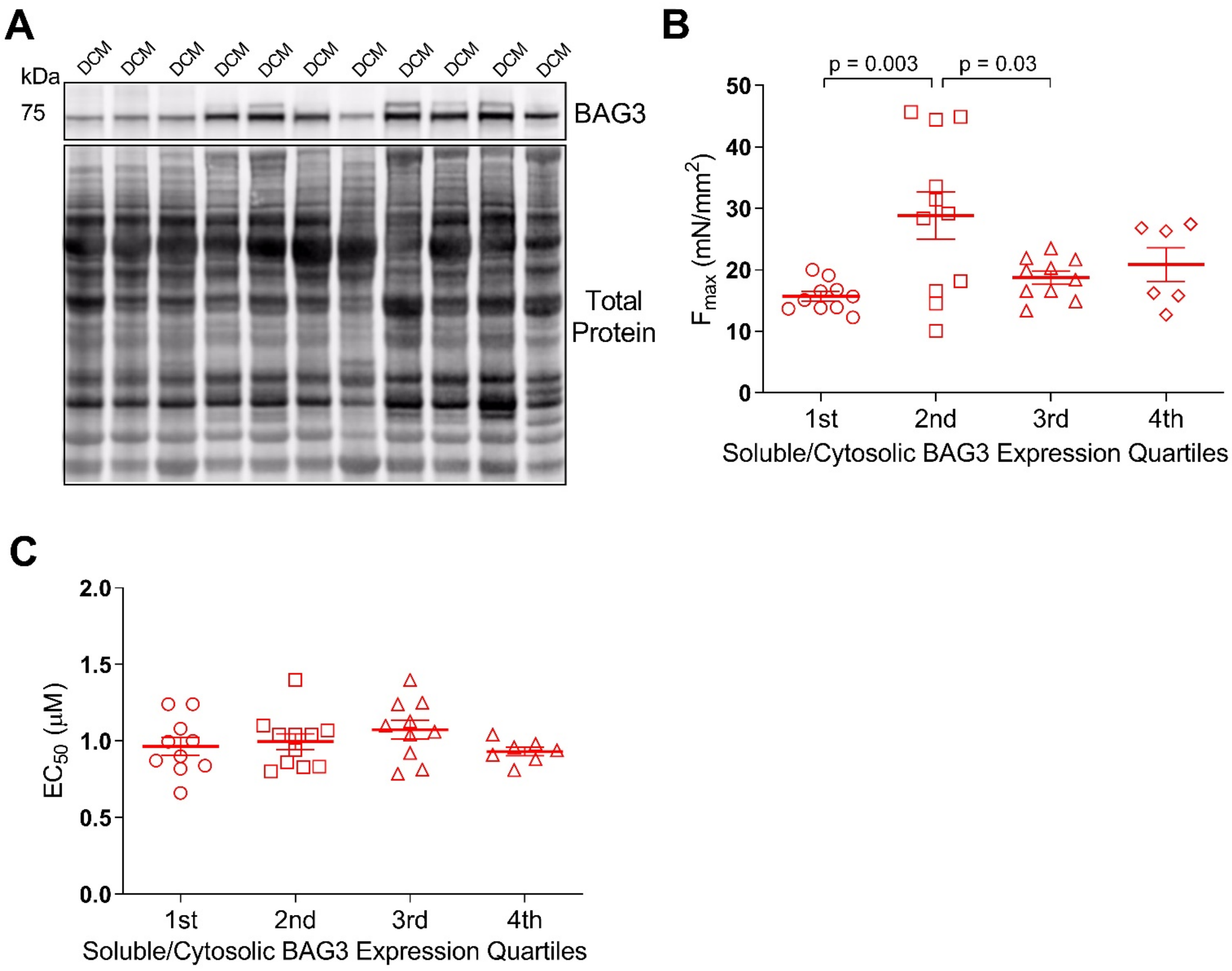


**Figure S1. Soluble/cytosolic BAG3 expression does not correlate with myofilament functional parameters. A**. Western blot for BAG3 in the triton-soluble fraction from the DCM LV samples. **B-C**. Myocyte $F_{max}$ (B) and $EC_{50}$ (C) values in the DCM samples organized by quartile of soluble BAG3 expression; $1^{st}$ = lowest BAG3 expressors, $4^{th}$ = highest; n = 11 DCM samples (one sample was not prepped for the soluble fraction due to lack of remaining tissue), 3-4 myocytes per sample for functional assessment. Data are presented as mean ± SEM and were analyzed using one-way ANOVA, Tukey post-hoc.

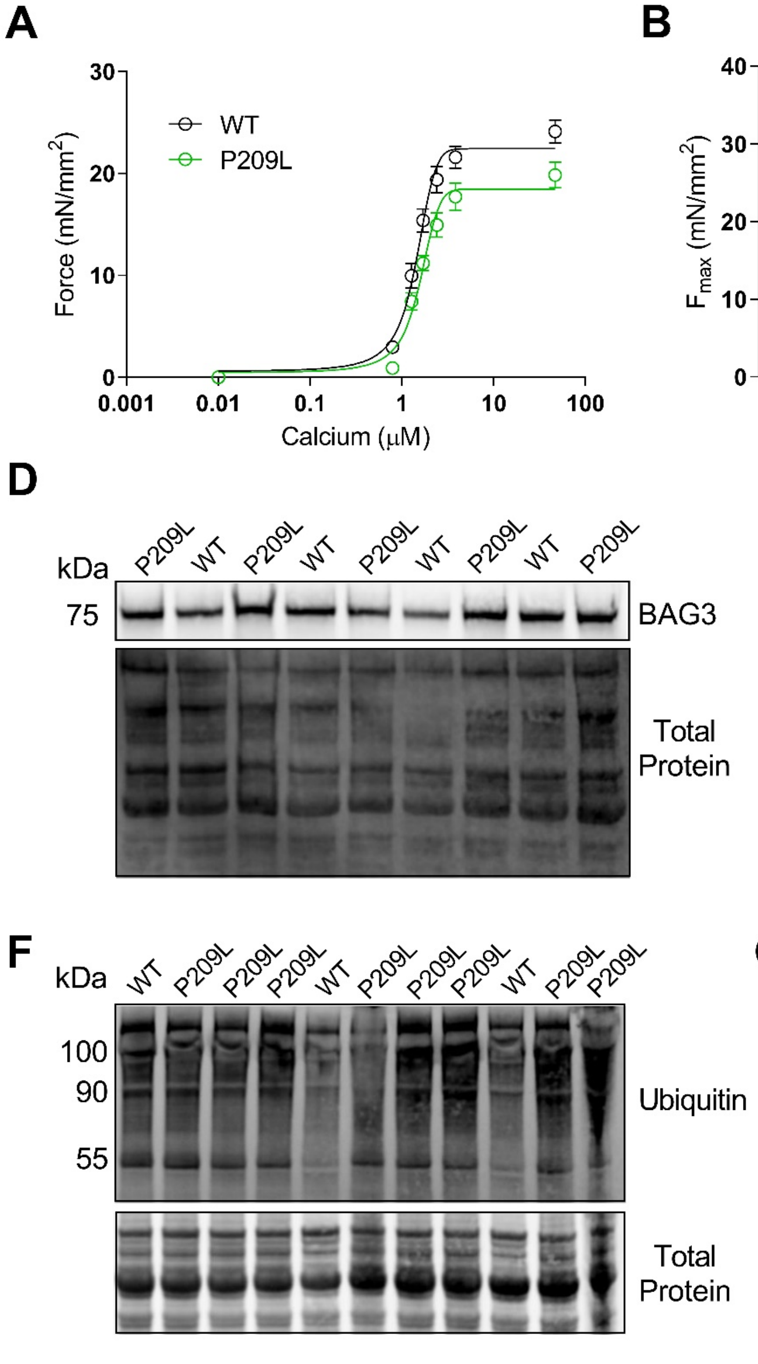


**Figure S2. The P209L BAG3 mutation decreases sarcomere contractile force and impairs myofilament protein turnover. A.** Skinned myocyte force-$Ca^{2+}$ relationship for wild-type and P209L myocytes. **B-C.** Summary data for individual myocyte $F_{max}$ (B) and calcium sensitivity (C); n = 18 WT myocytes from 6 mice, 15 P209L from 5 mice. **D-E.** Representative western blot for myofilament BAG3 (D) and normalization to total protein loading control (E) for the WT and P209L mice; n = 10 WT, 14 P209L. **F-G.** Representative western blot for myofilament ubiquitin (F) and normalization to total protein loading control (H) for the WT and P209L mice; n = 12 WT, 22 P209L. All data are presented as mean ± SEM ad were analyzed by 2-tailed *t*-test.

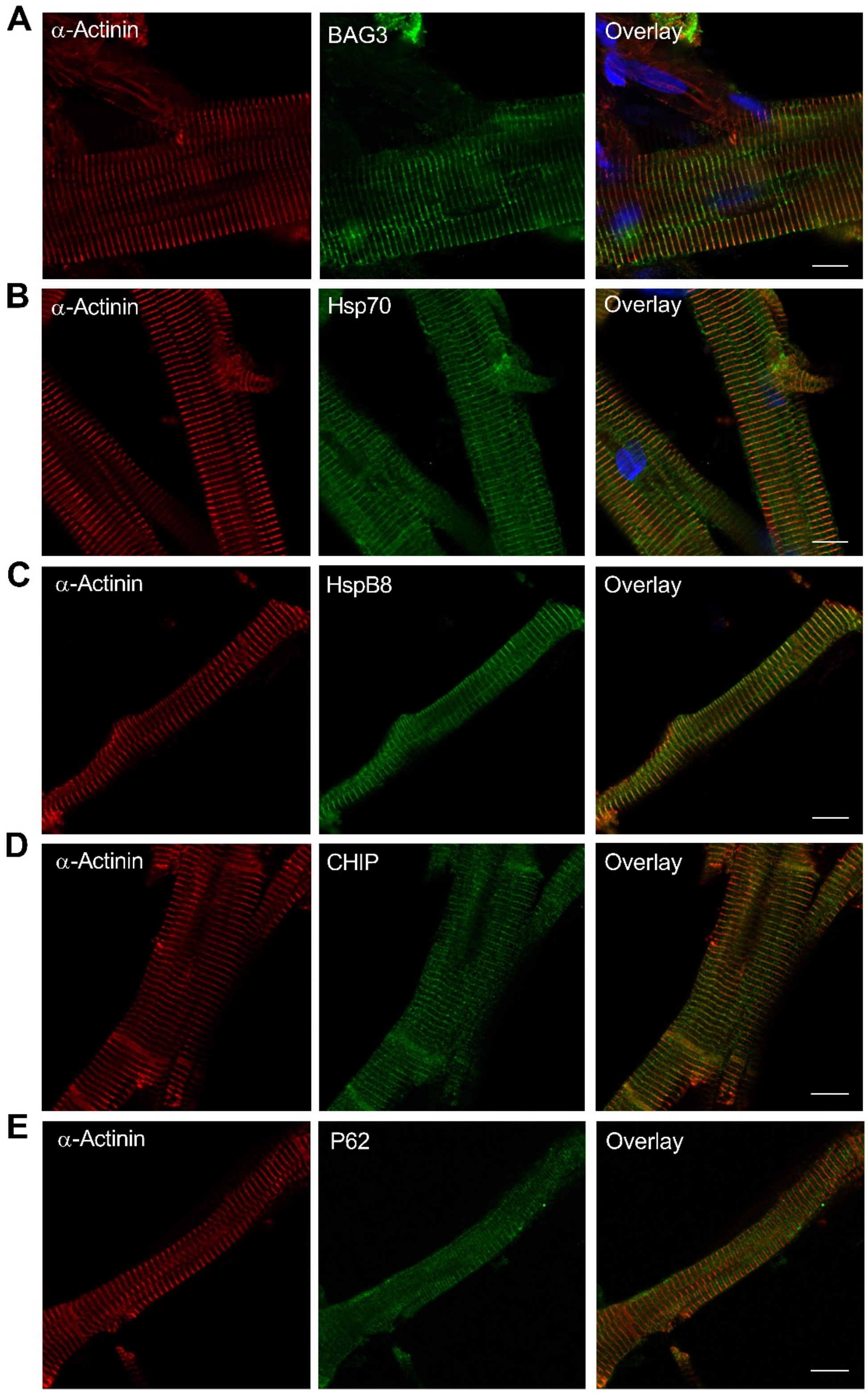


**Figure S3. Uncropped immunofluorescence images from the human LV myocyte experiments.** All images were acquired at 63X magnification; scale bars represent 10 microns.

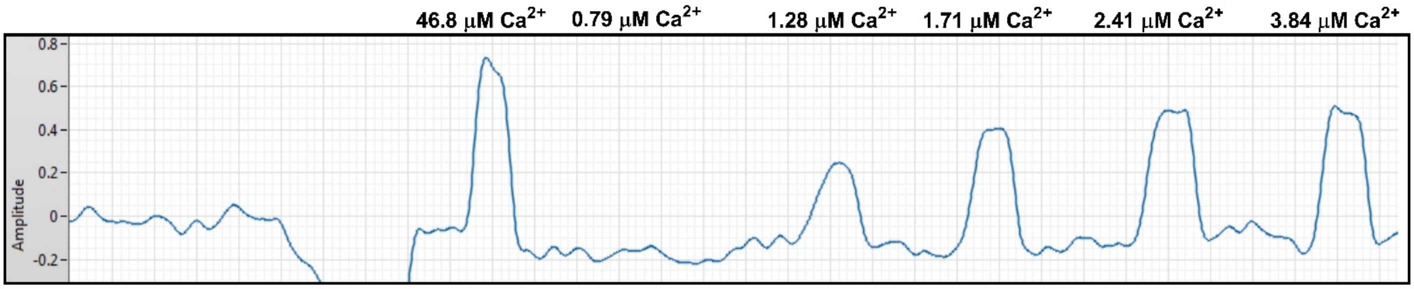


**Figure S4. Example recording from the skinned myocyte force-calcium experiments.** Single myocyte contractile force in response to various concentrations of calcium. Y-axis amplitude = millivolts; converted to mN using equation derived from force transducer calibration. Contractile forces were normalized to myocyte cross-sectional area during analysis.
